# Ultra-lightweight living structural material for enhanced stiffness and environmental sensing

**DOI:** 10.1101/2021.07.19.452887

**Authors:** Heechul Park, Alan F. Schwartzman, Tzu-Chieh Tang, Lei Wang, Timothy K. Lu

**Affiliations:** Synthetic Biology Group, Research Laboratory of Electronics, Massachusetts Institute of Technology, Cambridge, Massachusetts 02139, USA; Synthetic Biology Center, Department of Biological Engineering, Massachusetts Institute of Technology, Cambridge, Massachusetts 02139, USA; Department of Electrical Engineering and Computer Science, Massachusetts Institute of Technology, Cambridge, Massachusetts 02139, USA; Department of Materials Science and Engineering, Massachusetts Institute of Technology, Cambridge, Massachusetts 02139, USA

## Abstract

Natural materials such as bone, wood, and bamboo can inspire the fabrication of stiff, lightweight structural materials. Biofilms are one of the most dominant forms of life in nature. However, little is known about their physical properties as a structural material. Here we report an *Escherichia coli* biofilm having a Young’s modulus close to 10 GPa with ultra-low density, indicating a high-performance structural material. The mechanical and structural characterization of the biofilm and its components illuminates its adaptable bottom-up design, consisting of lightweight microscale cells covered by a dense network of amyloid nanofibrils on the surface. We engineered *E. coli* such that 1) carbon nanotubes assembled on the biofilm, enhancing its stiffness to over 30 GPa, or that 2) the biofilm sensitively detected heavy metal as an example of an environmental toxin. These demonstrations offer new opportunities for developing responsive living structural materials to serve many real-world applications.

---

Devising materials that are both lightweight and stiff is one of the challenges in structural materials because stiffer materials are generally heavier^1^. Such materials, however, have many uses in the aerospace and automobile industries and in the construction of portable devices, as well as other applications. Architectural designs in natural cellular materials such as bone, wood, and bamboo have inspired engineers to fabricate lightweight structural materials^2,3^. The properties of bacterial biofilms are also being explored, as these materials may offer not only principles of design but also the quality of self-renewal^4,5^. Biofilms are defined as interface-adherent multicellular communities encased in a self-produced extracellular matrix composed of polysaccharides, proteins, lipids, and extracellular DNA^4,6–8^. Although biofilms are often associated with recalcitrant infections arising in human medical and industrial settings^9^, they are indispensable to global biogeochemical cycling processes in nature^10^. Applications of biofilms have been demonstrated for the degradation of wastewater^11^, the monitoring of heavy metal pollutants in water^12^, and biocatalysis to produce chemicals and biofuels^13^. The viscoelastic properties of biofilms have also been extensively studied^14–17^, and biofilms have been shown as functional materials that can be engineered by synthetic biology^18–20^. However, little is known about their other physical properties, for example, their pure elasticity with density as a structural material. Moreover, current methods of monitoring heavy metals involve sampling biofilms and doing a cumbersome analysis of the metals absorbed into the biofilms^12^ rather than using programmable sensors and simple readouts. Biofilms have not been thoroughly explored and so remain largely unexploited as useful materials.

Here, we report a highly stiff *E. coli* biofilm with ultra-low density, indicative of a structural material having high performance capabilities compared to other natural and synthetic materials. We probe the biofilm and its structural components, which reveal a bottom-up design concept from the nanoscale to the macroscale, adaptable to building ultra-lightweight structural materials. Along with these findings, we demonstrate that, by interfacing this biofilm with inorganic nanomaterials, its elastic properties can be significantly enhanced. We also demonstrate that the biofilm can be transformed, by engineering, into a programmable living structural material that senses and responds rapidly to environmental toxins such as heavy metal. We have taken a synthetic biology approach to engineer this unusual material and provide a proof-of-concept for its development for real-world use.

## Structural and elastic properties of *E. coli* biofilms

To test the stiffness of the *E. coli* biofilm by instrumented indentation, a densely packed structure is desirable. We used *E. coli* MG1655 *PRO* Δ*csgA ompR234* (*E. coli* ompR234) having an integrated synthetic riboregulator producing CsgA amyloid fibrils, called curli (Supplementary Tables 1, 2). This choice of strain was based on: its substantial synthesis of self-assembled CsgA fibrils^21^; its lack of colonic acid synthesis, such that the biofilm it produces is non-mucoid^22^; and its fibril production, which is controllable by a chemical inducer, anhydrotetracycline (aTc). The addition of aTc induces the transcription of not only csgA mRNA but also *trans*-activating RNA (taRNA), preventing the cis-repressive (cr) sequence from blocking the ribosome-binding sequence (RBS) (Fig. 1a). When an increased seed concentration of *E. coli* ompR234 cells was used to grow the biofilm in a static medium for 44 hours, a closely packed self-organized structure was formed (Fig. 1b and Supplementary Fig. 1). Thioflavin T staining confirmed the production of amyloid curli fibrils from *E. coli* ompR234 (Supplementary Fig. 2). The thickness and biomass of the biofilm were analyzed from the cells constitutively expressing the red fluorescent protein mCherry by confocal microscopy and found to be 26.52 ± 4.59 *μ*m and 19.02 ± 4.10 µm^3^/µm^2^, respectively (Fig. 1b). The root-mean-square surface roughness of the biofilm over 2 *μ*m × 2 *μ*m areas was 32.3 ± 5.0 nm, determined by analysis of atomic force microscopy (AFM) height images (Supplementary Fig. 3).

**Fig. 1.**
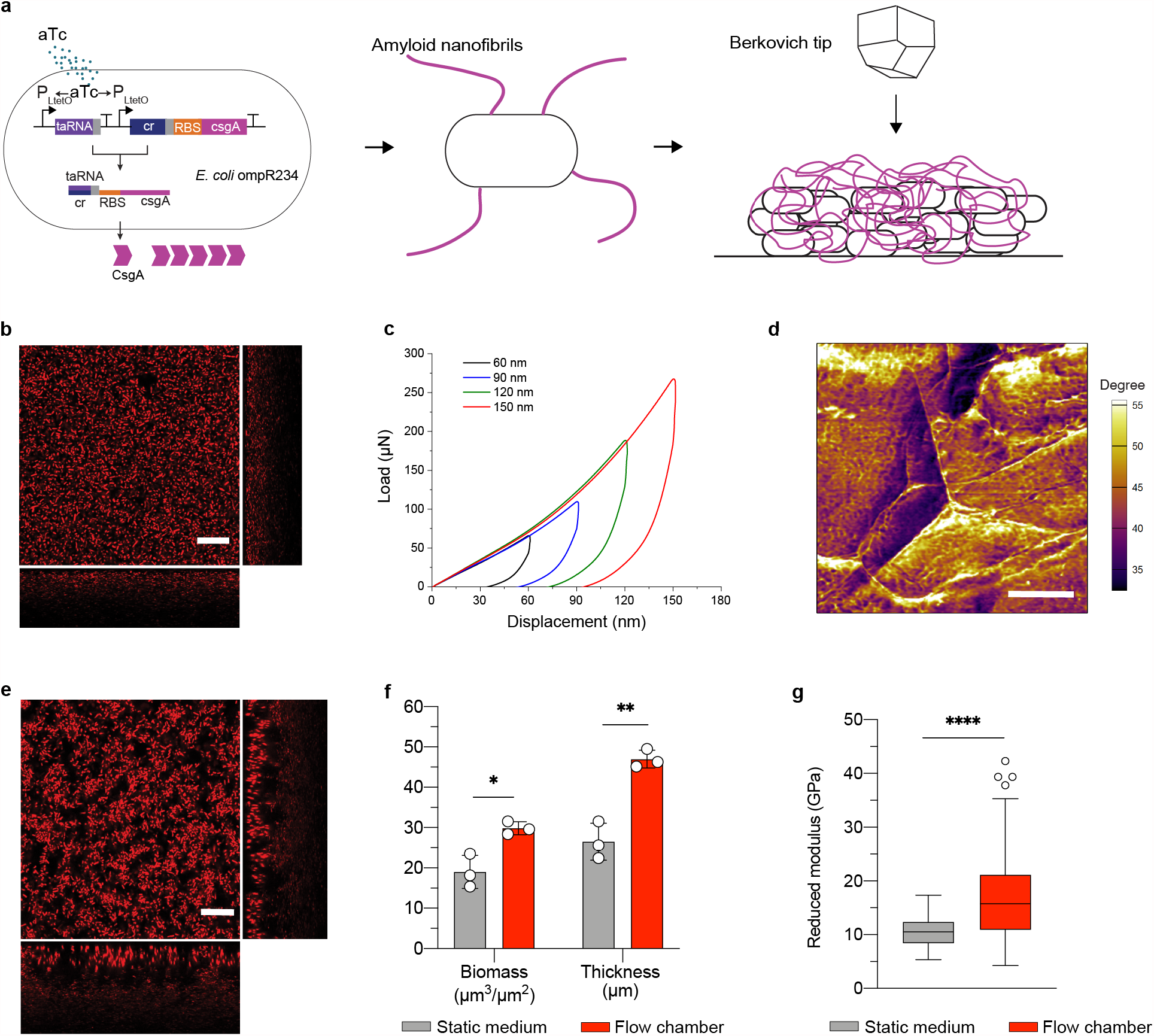
Structural and elastic properties of *E. coli* ompR234 biofilms grown in static medium and flow chamber. **a**, Schematic of a biofilm with inducible CsgA curli nanofibrils and stiffness characterization with instrumented indentation using a Berkovich probe tip. **b, e**, Confocal microscopy images with orthogonal XZ and YZ views of biofilms grown **b**, in a static medium and **e**, in a flow chamber. Scale bar, 20 *μ*m. **c**, Lines denote mean load-displacement curves on the static medium biofilm with multiple indenting depths of 60, 90, 120, and 150 nm as characterized by instrumented indentation. **d**, Atomic force microscopy (AFM) phase image showing an indented area after a 150 nm instrumented indentation was made in the static medium biofilm. Scale bar, 500 nm. **f, g**, Comparison between static medium and flow chamber biofilms **f**, for biomass and thickness and **g**, for reduced modulus. **f**, Data are mean ± s.d. for n = 3 independent samples. **g**, Box-plots show center lines (median), box limits (upper and lower quartiles), whiskers (1.5× interquartile range), and points (outliers) for n ≥ 140 from three independent samples. **f, g**, *P≤0.05, **P≤0.01, ****P≤0.0001, Student t test.

To characterize the elastic modulus of the *E. coli* ompR234 biofilm, we applied instrumented indentation using a Berkovich diamond probe tip (Fig. 1a). All indentations in this study were carried out in displacement-controlled mode with a displacement of 150 nm, unless otherwise noted. A trapezoidal load function was used to minimize the nose effect upon unloading due to creep on load-displacement curves. The instrumented indentation measurements revealed that the *E. coli* ompR234 biofilm had a reduced elastic modulus of 10.5 ± 2.6 GPa, as analyzed by the Oliver-Pharr method^23^ (Fig. 1c). The indentation results for other displacements, 60, 90, and 120 nm, yielded similar moduli (Fig. 1c and Supplementary Fig. 4), which ruled out the possibility of a substrate effect. Moreover, as imaged by the AFM tapping mode, a contact impression remained after a 150 nm-displacement indentation (Fig. 1d), confirming that the biofilm had been clearly indented by the Berkovich tip. This AFM contact impression and the residual displacements observed in the load-displacement curves (Fig. 1c, d) also indicate the plastic nature of the indentation in the biofilm. The Young’s modulus of the biofilm was 9.6 ± 2.4 GPa, as calculated from the reduced modulus and assuming a Poisson’s ratio of 0.3. This Young’s modulus indicates that the biofilm formed by *E. coli* ompR234 is exceptionally stiff compared to other biofilms, such as those of *P. aeruginosa* (30 Pa-20 kPa)^24–26^ and *B. subtilis* (1-50 kPa)^27,28^.

Because flow can affect structural and mechanical properties^7^, the *E. coli* ompR234 biofilm was also grown in a microfluidic flow chamber, with dimensions of 250 *μ*m high × 2 mm wide × 20 mm long (Supplementary Fig. 5). To grow a densely packed biofilm, a bacterial suspension with the same seed concentration was continuously introduced into the microfluidic chamber for 44 hours. The confocal microscopy imaging showed a densely packed biofilm with a biomass of 29.83 ± 1.61 µm^3^/µm^2^ and a thickness of 46.99 ± 2.25 µm (Fig. 1e). The biomass and thickness increased by 57% and 77%, respectively, compared to those of the static medium biofilm (Fig. 1f). The flow chamber biofilm measured by instrumented indentation showed a reduced elastic modulus of 16.7 ± 7.7 GPa, which is an increase of 62 % compared to that of the static medium biofilm (Fig. 1g). The Young’s modulus was 15.4 ± 7.1 GPa, as calculated from the reduced modulus and assuming a Poisson’s ratio of 0.3. Both the static medium and flow chamber biofilms are less stiff than a ceramic alumina material (393 GPa) or steel (200 GPa) but stiffer than a polystyrene plastic (3.5 GPa)^1^.

## Specific stiffness

Specific stiffness normalized by density is a standard property of materials, indicative of both stiffness and weight; higher values mean higher stiffness with lighter weight. To obtain the specific stiffness of the biofilm required determining its density. To determine a wet density, we investigated the mass of the static medium biofilm by using thermogravimetric analysis (TGA). The TGA results revealed the area density to be 105.8 ± 8.1 µg/cm^2^ at 37°C (Fig. 2a). The biofilm’s volume was obtained from the biomass (by definition, biomass is the volume of all voxels that contain biological entities divided by the substrate area^29^). By dividing the area density by the biomass, the density was estimated to be 0.057 ± 0.011 g/cm^3^. The biofilm is thus ultra-lightweight compared to other stiff materials such as alumina (3.80 g/cm^3^) and steel (8.05 g/cm^3^), and over 18 times lighter than polystyrene (1.05 g/cm^3^)^1^. By dividing the Young’s modulus by the density, we determined the specific stiffness of the *E. coli* ompR234 biofilm to be 173.8 ± 34.5 GPa/g·cm^-3^. This value was plotted in an Ashby plot^3^ for comparison (Fig. 2b). The biofilm thus outperforms the other natural materials that have been characterized so far, such as bone, shell, and wood, and its property of specific stiffness surpasses even that of most metallic alloys. The *E. coli* ompR234 biofilm, therefore, is comparable to high-performance materials such as silicon carbide, which has a specific stiffness of 140 GPa/g·cm^-3^, and boron carbide, which has a specific stiffness of 182 GPa/g·cm^-3^ (see ref.^1^).

**Fig. 2.**
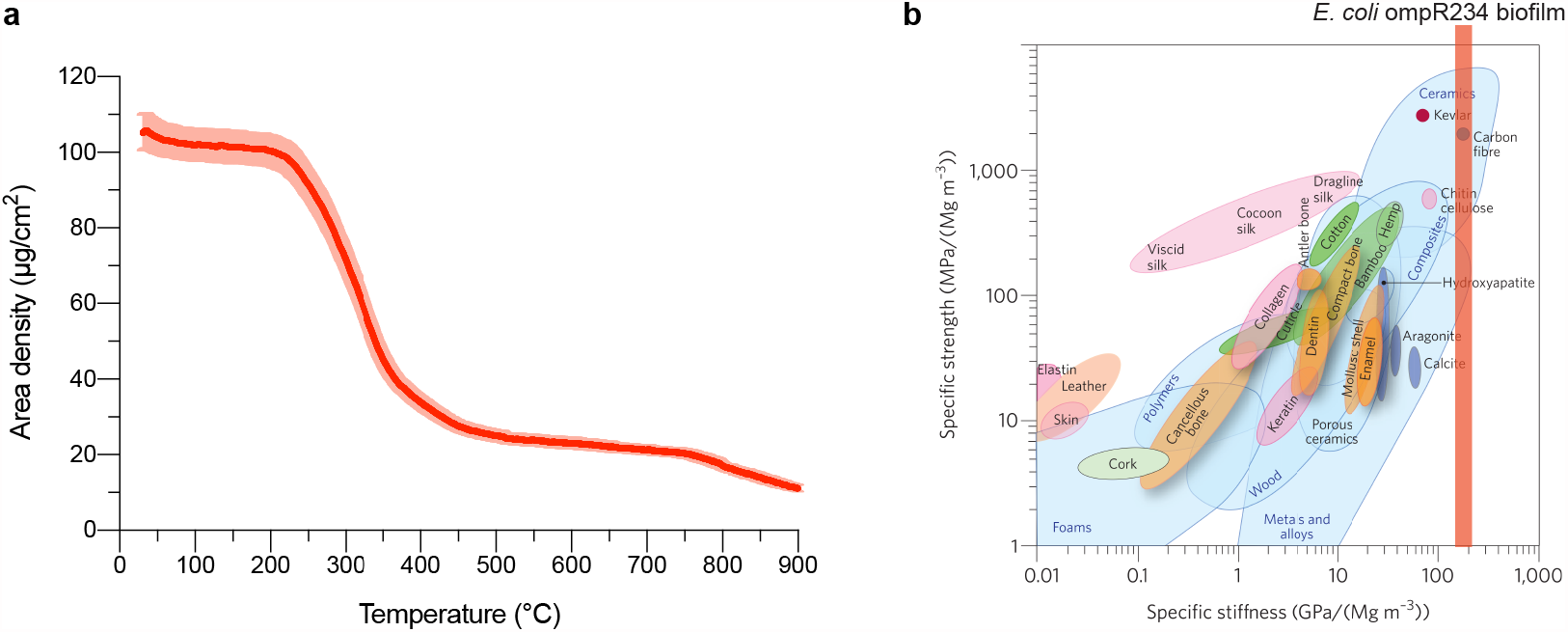
Area density and specific stiffness of the *E. coli* ompR234 biofilm. **a**, The red line denotes a mean curve of area density as a function of temperature by thermogravimetric analysis (TGA). The mass and volume of the biofilm were obtained from this area density curve and the biomass, respectively, to estimate its density. Data are plotted as mean ± s.d. for n = 3 independent samples. **b**, The specific stiffness of the biofilm was plotted in an Ashby plot^3^. The panel **b** was reprinted by permission from Springer nature customer service centre GmbH : Springer nature, Nature materials, Bioinspired structural materials, Wegst, U. G. K. *et al*., copyright 2015.

## Nanofibrils and cells, and dose effect

To decouple the structural components of the biofilm, we independently probed the amyloid fibrils and the *E. coli* ompR234 cells. The self-assembled fibrils, one of the stiffest protein materials, are a key determinant of the structural properties of the *E. coli* ompR234 biofilm^22,30^. The CsgA amyloid fibrils had a radial Young’s modulus of 1.92 ± 0.72 GPa, as determined by AFM quantitative mapping of elasticity (Fig. 3a, b, c). This gigapascal level stiffness apparently originates from the characteristic cross-β structure of the amyloid fibrils, which have dense hydrogen-bonding networks^31^. The Young’s modulus of the *E. coli* ompR234 cells was 24.2 ± 6.8 MPa, probed by AFM force spectroscopy (Fig. 3d, e, f). Thus, the cells are much less stiff than the CsgA amyloid fibrils. As far as the dimensions of each component, the rod-shaped cells were 0.56 ± 0.04 *μ*m in width and 1.91 ± 0.42 *μ*m in length, as analyzed from scanning electron microscope (SEM) images (Supplementary Fig. 6). The curli fibrils have been reported to be approximately 4-11 nm in width and on the order of micrometers in length^32^. The SEM image shows that a dense network of amyloid nanofibrils not only covers the cell surface but also occupies the spaces between the cells on the biofilm surface (Fig. 3g). Thus, the biofilm’s ultra-low density comes from the bulk volume of the microscale cells while the high stiffness stems from the dense nanofibril network on the surface. To make a less dense fibril network, we investigated the dose effect of lower concentrations of the inducer aTc on the static medium biofilm (Supplementary Fig. 7). Compared to the original aTc concentration of 250 ng/mL, the thickness, biomass, and modulus at 12.5 ng/mL aTc significantly decreased by 31%, 51%, and 51%, respectively (Fig. 3h, i). Because the biomass analyzed in this research also indicates the approximate biomass of the nanofibrils (Supplementary Fig. 8), the decreased modulus observed at 12.5 ng/mL aTc was understood to result from a decreased biomass of the nanofibrils. In addition, the increased modulus of the flow chamber biofilm over that of the static medium biofilm was due to an increased biomass of the nanofibrils (Fig. 1f, g).

**Fig. 3.**
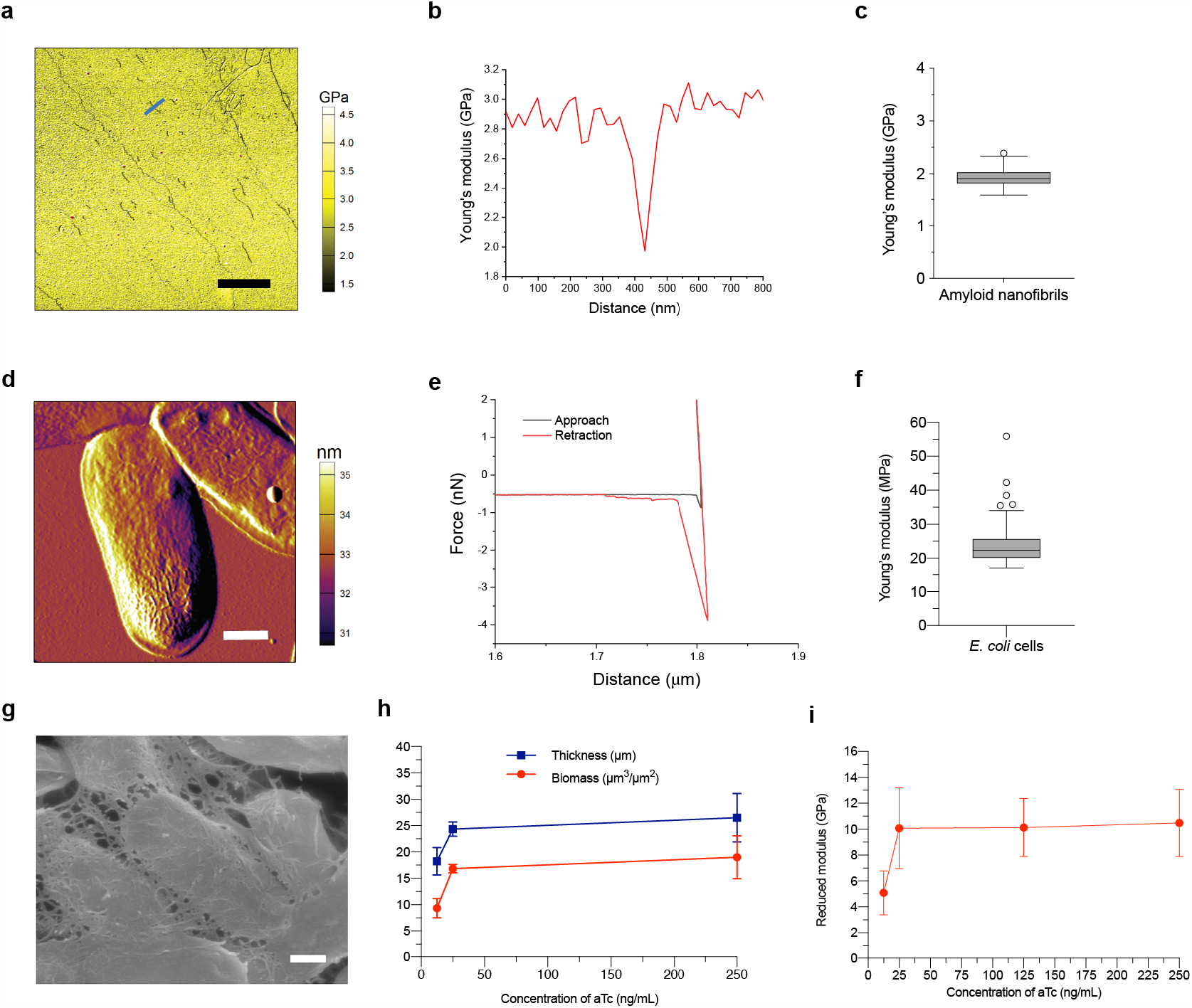
Elastic properties of the *E. coli* ompR234 biofilm’s components and dose effect of an inducer on the biofilm. **a**, AFM image of elastic mapping of the nanofibrils. The nanofibrils were prepared on a polystyrene substrate with a known elastic modulus of 2.7 GPa. Scale bar, 2 µm. **b**, Representative cross-section analysis of an amyloid nanofibril indicated by the blue line in **a** with the polystyrene background. **c**, The cross-section analysis resulted in radial Young’s moduli of the nanofibrils. **d**, Amplitude image of AFM tapping mode on *E. coli* ompR234 cells. Scale bar, 500 nm. **e**, Representative force-distance curves of the cells for both approach and retraction in AFM contact mode force spectroscopy. **f**, The resulting Young’s moduli of the cells are shown. **c, f**, Box-plots show center lines (median), box limits (upper and lower quartiles), whiskers (1.5× interquartile range), and points (outliers) **c**, for n = 77 from three independent samples and **f**, for n = 64 from four independent samples. **g**, Scanning electron microscope (SEM) image of the biofilm surface. Scale bar, 300 nm. **h**, Biomass and thickness of the biofilm as a function of the concentration of an inducer, anhydrotetracycline (aTc). **i**, Reduced elastic moduli of the biofilm as a function of aTc concentration. **h, i**, Data are plotted as mean ± s.d. **h**, for n = 3 independent samples and **i**, for n ≥ 125 from three independent samples.

## Enhanced stiffness by carbon nanotubes

To determine whether the *E. coli* ompR234 biofilm could be engineered to enhance its elastic properties by interfacing it with inorganic nanomaterials, we chose single-walled carbon nanotubes (SWNTs) (Fig. 4a). SWNTs show a high aspect ratio of 1 nm in diameter and 500 nm in length and a high Young’s modulus^33^. To bind SWNTs to the nanofibril network, we used the short peptide SpyTag and its cognate protein SpyCatcher, which form a covalent amide bond between them^34^. The *E. coli* ompR234 strain was engineered to display SpyTag at the C-terminus of the nanofibril, inducible by aTc (Supplementary Tables 1, 2). The purified SpyCatcher proteins used here have two cysteine and six histidine residues at the N-terminus (Supplementary Fig. 9). SWNTs were dispersed in surfactant-dissolved solution via sonication and purified via centrifugation (Fig. 4b). The purified SWNTs were then non-covalently functionalized with SpyCatcher (Fig. 4c). The non-covalent functionalization of the SWNTs arises from the binding affinity of the six aromatic histidine residues on SpyCatcher for the SWNT through π-π stacking^35^. The SpyCatcher-to-SWNT ratio of 1,000:1 was maximally dispersed, while SWNTs without SpyCatcher aggregated (Fig. 4d). Higher photoluminescence intensity suggests better dispersion of SWNTs, because metallic SWNTs in small bundles quench photoluminescence^36^. To reinforce the biofilm, the SpyCatcher-functionalized SWNTs (1,000:1 SpyCatcher-to-SWNT ratio) were assembled on the SpyTag nanofibril network. The SWNT-assembled biofilm was then compared to a bare biofilm including the SpyTag amyloid nanofibrils but without SWNTs. The transmission electron microscopy images of the bare SpyTag nanofibrils and SWNT-assembled nanofibrils scraped from the two kinds of the biofilms (Fig. 4e, f) confirmed the integration of the SWNTs into the biofilm. Instrumented indentation showed that the SWNT-assembled biofilms had reduced elastic moduli of up to 32.3 ± 11.7 GPa, which was 3.2 times the stiffness of the bare SpyTag biofilm of 10.2 ± 3.6 GPa (Fig. 4g). The 32.3 GPa reduced modulus corresponds to a Young’s modulus of 30.7 ± 10.7 GPa, assuming a Poisson’s ratio of 0.3.

**Fig. 4.**
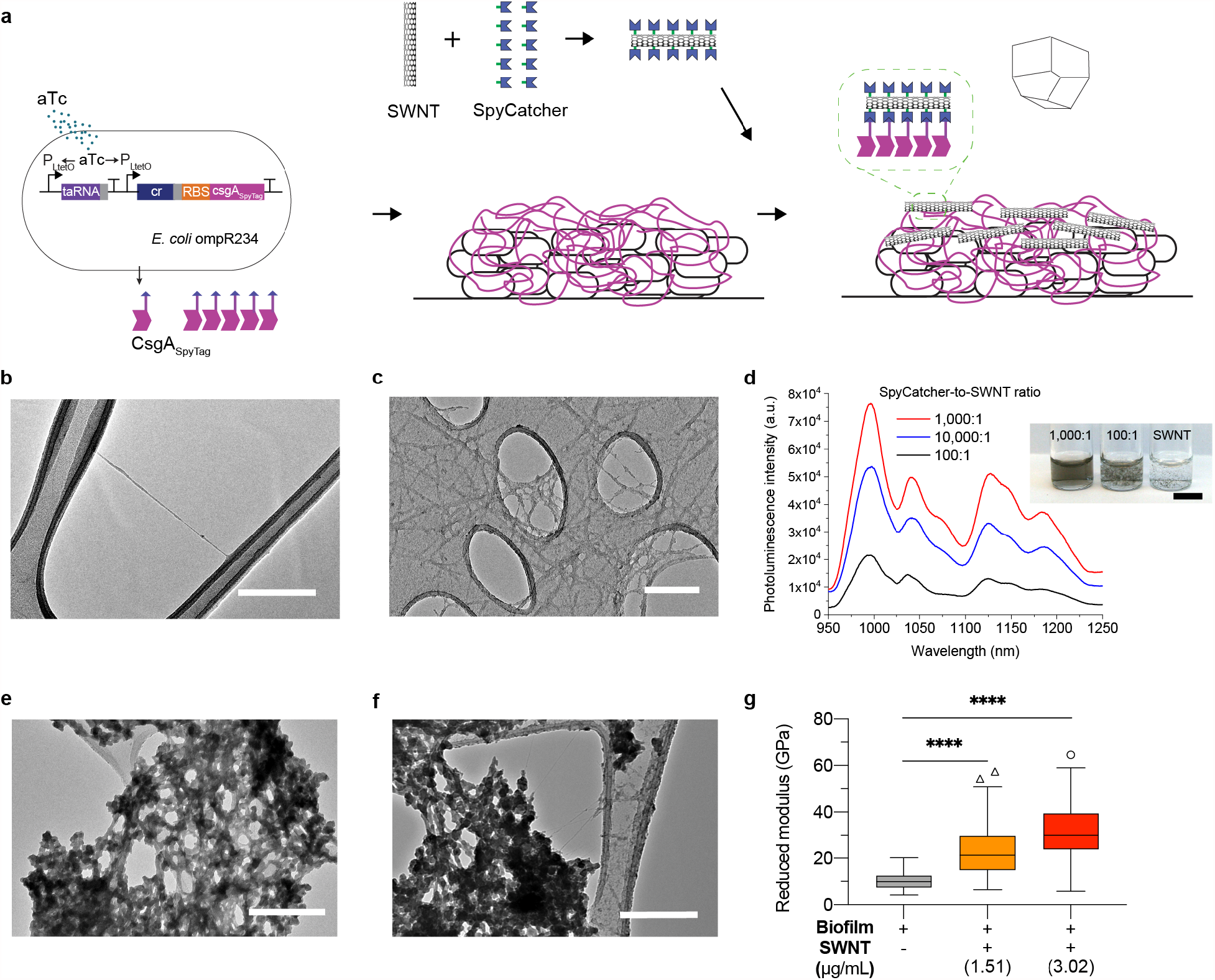
Enhancement of biofilm stiffness by single-walled carbon nanotubes (SWNTs). **a**, Schematic of binding SWNTs to a nanofibril network on a biofilm. **b**, Transmission electron microscopy (TEM) image of a purified SWNT dispersed in sodium cholate solution. Scale bar, 200 nm. **c**, TEM image of SWNTs functionalized with SpyCatcher (1,000:1 SpyCatcher-to-SWNT ratio). Scale bar, 200 nm. **d**, Photoluminescence excited at 785 nm at the same absorbance at 632 nm. The inset is a photograph taken under ambient light. Scale bar, 0.5 cm. **e**, TEM image of SpyTag nanofibrils scraped from the bare biofilm. Scale bar, 200 nm. **f**, TEM image of SWNT-assembled SpyTag nanofibrils scraped from the biofilm after binding SWNTs. Scale bar, 200 nm. **g**, Stiffness results of instrumented indentation. The concentration (µg/mL) indicates the seed density of the functionalized SWNTs. Box-plots show center lines (median), box limits (upper and lower quartiles), whiskers (1.5× interquartile range), and points (outliers) for n ≥ 145 from three independent samples. ****P≤0.0001, Student t test.

## Responsive living material detecting cadmium

To engineer the biofilm to act as an environmental living sensor, we integrated an additional metal-sensing plasmid into *E. coli* ompR234 to express green fluorescent protein (GFP) upon the detection of heavy metals (Fig. 5a and Supplementary Tables 1, 2). The metal-sensing plasmid was composed of ZntR, a transcriptional regulator activated by metal ions (Cd^2+^, Pb^2+^, or Zn^2+^), and the promoter PzntA to express GFP^37^. The resultant biofilm was tested for both kinetic and end-point responses at room temperature in M63 medium for the detection of cadmium as an example of heavy metals. The biofilm responded rapidly to cadmium ions in 20 minutes and the kinetic response became saturated at around 9 hours (Fig. 5b). To activate the entire quantity of sensor cells for the end-point measurements, the biofilm was incubated for 9 hours in medium containing 0, 0.05, 0.5, or 5 ppm cadmium. The results indicate that the living biofilm material had sufficient sensitivity to detect cadmium ions at a concentration as low as 0.05 ppm (Fig. 5c). Thus, the living biofilm material exhibited the sensitive, rapid detection of cadmium. This demonstration suggests that biofilms can be transformed into highly responsive living structural materials with genetic sensors for a wide variety of sensing applications.

**Fig. 5.**
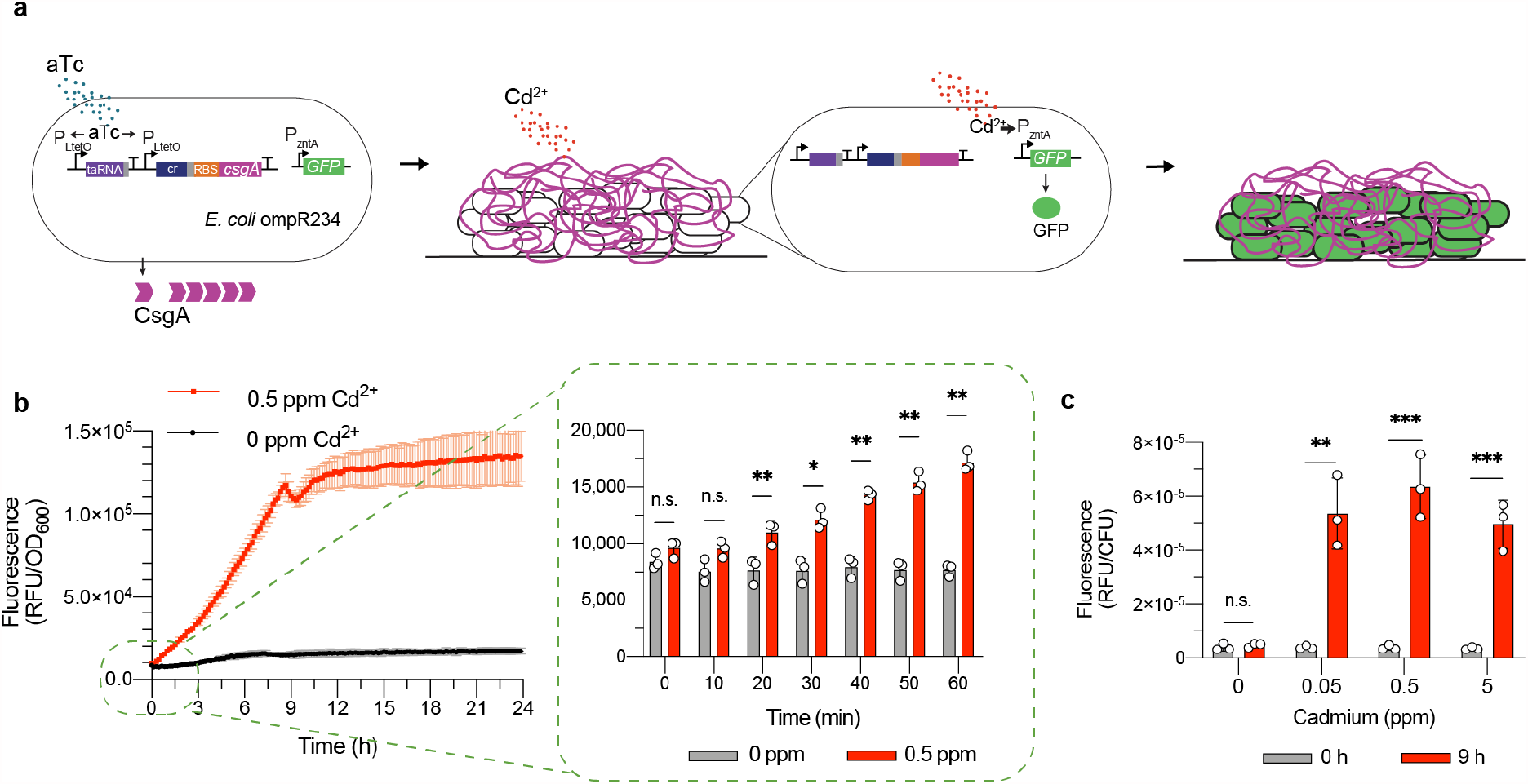
Living biofilm material to sense and respond to environmental toxins. **a**, Schematic of a metal-sensing biofilm material. **b**, Kinetic response of the engineered living biofilm material detecting cadmium. The inset shows a short timescale of 0 – 60 minutes. **c**, Results of the GFP fluorescence measurements at 0 and 9 hours of exposure to cadmium. **b, c**, Data are mean ± s.d. for n = 3 independent samples. Not significant (n.s.) P>0.05, **P≤0.01, ***P≤0.001, Student t test. GFP, green fluorescent protein; RFU, relative fluorescence unit; OD_600_, optical density at 600 nm; CFU, colony-forming unit.

## Conclusions

Biofilms are among the most successful and abundant modes of life on Earth^6^ and, along with bone, wood, and bamboo, constitute valuable structural materials. Biofilms can be exploited not only indirectly as designs that can inspire the development of new synthetic materials but also directly as practical materials. We demonstrated that the *E. coli* ompR234 biofilm has a high Young’s modulus, close to 10 GPa, with ultra-low density. The resulting specific stiffness of over 170 GPa/g·cm^-3^ indicates that the *E. coli* ompR234 biofilm is a high-performance structural material comparable to other structural materials such as silicon carbide and boron carbide. We also showed that by assembling single-walled carbon nanotubes on its surface, the biofilm could be engineered to increase its elastic moduli up to over 30 GPa.

Investigating the mechanical and structural properties of the amyloid nanofibrils, cells, and biofilm suggests an important bottom-up design concept for creating lightweight structural materials: soft lightweight materials of bulk volume can be significantly stiffened by covering them with a strong dense fiber network. In the specific case shown here, microscale cells are the relatively soft bulky material that is covered and reinforced by a dense network of nanoscale amyloid fibrils. This arrangement, along with the ability of the biofilm to self-organize, results in a macroscale high-performance material. The dense nanofibril network on the biofilm surface is a distinct property of the *E. coli* ompR234 biofilm. This biofilm has higher moduli than the *P. aeruginosa* and *B. subtilis* biofilms described so far, which have polysaccharides^26,38^ and BslA hydrophobins^27,39^ on their surface, respectively, that contribute to their much lower moduli.

This study also demonstrated that the high-performance biofilm characterized here can be transformed into highly responsive living structural materials that sense and respond to specific substances for useful industrial applications. As biofilms are globally abundant, we envision that they can be easily deployed to act as inexpensive environmental sensors to alert people to the presence of toxins. Examples include heavy metal sensors to detect contaminants on a bathroom washbasin and steroid hormone sensors^40^ on a rock in a stream for monitoring water quality. We also envision that, owing to their surface-adhesive property, high-performance biofilms can be used as coating materials on portable products and textiles^41^, to sense environmental toxins. We anticipate that as the development of biofilm-based sensors detecting environmentally relevant toxins proceeds, the range of potential uses for them will expand.

## Methods

### Strains and seed cultures

All strains and plasmids used in this study are listed in Supplementary Tables 1 and 2. For instrumented indentation and AFM characterizations, the *E. coli* MG1655 *PRO* Δ*csgA ompR234* strain with the synthetic riboregulator was used to tightly regulate the expression of CsgA amyloid fibrils. The synthetic riboregulator is responsive to the chemical inducer anhydrotetracycline (aTc). Thus, CsgA expression was controlled by aTc. For the SWNT experiment, the *E. coli* strain with the synthetic riboregulator was engineered to display SpyTag on the CsgA fibrils, as follows. DNA sequences encoding the tag were inserted at the 3’ end of the aTc-inducible *csgA* gene. For confocal microscopy, the strain carrying both the synthetic riboregulator and a plasmid constitutively expressing a fluorescent protein (mCherry) was used. For sensing and responding to heavy metal ions, *E. coli* MG1655 *PRO* Δ*csgA ompR234* was transformed with the metal-sensing plasmid as well as the CsgA synthetic riboregulator. Seed cultures were grown in LB-Miller medium overnight at 37°C at 250 r.p.m with the following antibiotics: 25 µg/mL chloramphenicol, 100 µg/mL carbenicillin, 50 µg/mL spectinomycin, and 50 µg/mL kanamycin. The cells were then washed twice with phosphate-buffered saline (PBS).

### Biofilm growth in static medium

M63 minimal medium (Amresco, Inc.) supplemented with 0.2% w/v glucose and 1 mM MgSO_4_ was used for biofilm growth. To induce CsgA curli, 250 ng/mL aTc was added, along with the following antibiotics: 25 µg/mL chloramphenicol, 100 µg/mL carbenicillin, 50 µg/mL spectinomycin, and 50 µg/mL kanamycin. The supplemented M63 medium was inoculated at the seed density of 2.26 × 10^8^ cells/mL from the seed cultures. The biofilms were grown at 30°C for 44 hours in the medium and then washed twice with deionized water for characterization.

### Microfluidic flow chamber device fabrication

The single-channel microfluidic flow chambers, with dimensions of 250 *μ*m high × 2 mm wide × 20 mm long, were fabricated using a standard soft lithography with polydimethylsiloxane (PDMS)^42^. Patterns for the microfluidic chambers were designed using the software AutoCAD (Autodesk, Inc.). The chambers were printed on transparent films (Advance Reproduction Corporation, MA, USA) and used as a photomask. A mold (255 *μ*m high) was fabricated in two steps, according to the electronic visions model EV620 mask aligner (10 mW cm^−2^, EVGroup, Austria): first, the photoresist SU8-2005 was used to form a 5 *μ*m attaching layer between a silicon wafer and the top photoresist layer; then the top 250 *μ*m photoresist layer was made with SU8-2100. After the mold fabrication, a well-mixed and degassed PDMS prepolymer [RTV 615 A and B (10 : 1, w/w)] was poured onto the fluidic mold to yield a 3 mm-thick fluidic layer. Then, the mold was baked for 1 hour at 80°C. After baking, the PDMS layer structure was peeled from the mold. Two holes were punched by a metal biopsy punch (Robbins Instruments, Inc.) at the terminals of the chamber for an inlet and outlet. The PDMS chamber on a glass coverslip was gently pressed by two custom-made acrylic clamps to prevent leakage. A syringe was connected with silicone tubes to the inlet of the microfluidic chamber via a bubble trap (IBI Scientific).

### Biofilm growth in microfluidic chambers

The biofilms were grown on #1.5 rectangular glass coverslips, 22 × 40 mm (Ted Pella, Inc) by the flow of a bacterial suspension of 2.26 × 10^8^ cells/mL in the supplemented M63 medium. The suspension was introduced continuously by a syringe pump (Pump 11 Elite, Harvard Apparatus) into the microfluidic PDMS chamber at room temperature (24-26°C) for 44 hours at 1 *μ*L/min, which corresponds to an average flow speed of 33 *μ*m/s. After biofilm growth, deionized water was flushed through at 500 *μ*L/min for 40 seconds to remove planktonic cells and residual M63 medium.

### Scanning electron microscope (SEM)

The biofilms (static medium) were grown on #1.5 round glass coverslips, 1 cm diameter (Ted Pella, Inc) in a 24-well plate. The biofilms on the coverslips were fixed for 15 min with 4% paraformaldehyde and washed twice with deionized water. The biofilms were then dehydrated in a graded ethanol series of 30, 50, 70, 90, and 100%, followed by critical point drying (Tousimis autosamdri-815). After 2 nm Au/Pd sputter coating, the biofilms were imaged using a JEOL JSM 6700F SEM operated at 3-5 kV accelerating voltages.

### Confocal laser scanning microscopy (CLSM)

The biofilms (static medium) were grown in a 96-well plate with #1.5 glass bottom (MatTek Corporation). The biofilms (flow chamber) were grown on #1.5 rectangular glass coverslips, 22 × 40 mm (Ted Pella, Inc). CLSM was performed at room temperature (23°C) by using a Zeiss Laser Scanning Confocal Microscope LSM 710 with a 60x/1.40 oil Plan-Apochromat objective. For biomass and thickness, the bacterial cells of the biofilm constitutively expressing the red fluorescent protein mCherry were imaged in a single channel mode using Zen software (Zeiss). The excitation wavelength was 561 nm and the range of detection wavelengths was 585 - 665 nm. Z-stack images on the biofilms were collected over a scan area of 135 *μ*m × 135 *μ*m with an interval of 0.35 *μ*m, which guarantees at least 50% section overlaps. The Z-stack images were analyzed and the average biomass and thickness were obtained from at least three biofilms by the software COMSTAT^29^. The three-dimensional reconstruction of the Z-stack images was performed with Fiji, an ImageJ-software package. For confirmation of the amyloid fibrils, the fibrils in the biofilm were stained with a thioflavin T solution of 250 µg/mL (Sigma-Aldrich) for 5 min at room temperature. Then, the biofilms were additionally washed twice with deionized water. The image was obtained by confocal microscopy with excitation at 458 nm and detection from 465 - 535 nm.

### Thermogravimetric Analysis (TGA)

The mass of the biofilms was recorded on Discovery TGA (TA Instruments). The biofilms (static medium) were grown on 24-well plates and collected by pipetting 90 *μ*L deionized water per well. The collected samples were transferred to a platinum pan (TA instruments) that had been tared beforehand. To dehydrate solvent, the samples were incubated at 37°C for 4 hours inside a TGA furnace. After cooling down to 30°C, the samples were heated to 900°C at a heating rate of 10°C/min. Nitrogen gas was continuously injected at a rate of 25 mL/min throughout the TGA procedure. The average area density was obtained from three samples by dividing the measured mass by the substrate area.

### Instrumented indentation

The biofilms (static medium) were grown on #1.5 round glass coverslips, 1 cm diameter (Ted Pella, Inc) in a 24-well plate. The biofilms (flow chamber) were grown on #1.5 rectangular glass coverslips, 22 × 40 mm (Ted Pella, Inc). Both static and flow biofilms were then dried in air at room temperature (24-26°C) for 1 day before indentation. The Triboindenter 950 (Hysitron, Inc.) was used at room temperature (23°C) in ambient air for instrumented indentation. A trapezoidal load function was divided into three segments. For the 150 nm displacement, the first segment had a loading rate of 15 nm/s for 10 seconds. Once the maximum displacement was reached, the second segment of a hold period of 10 seconds followed. The third segment decreased the load at an unloading rate of 15 nm/s for 10 seconds. Multiple maximum displacements of 60, 90, and 120 nm with the same hold for 10 seconds and constant rates of loading and unloading were measured. More than 36 indents per sample were made in a square grid arrangement with each indent spaced 5 *μ*m apart for at least three biofilms. The Berkovich diamond area function and machine compliance were calibrated using a fused quartz substrate before each set of measurements. Reduced elastic moduli were extracted from initial unloading curves with an upper limit of 95% and lower limit of 40%, based on the Oliver-Pharr analysis^23^, using a Triboindenter software (Hysitron, Inc.). The reduced modulus was converted to Young’s modulus assuming Poisson’s ratio of 0.3 with the following equation: 1/E_r_ = (1-v_i_^2^)/E_i_ + (1-v_s_^2^)/E_s_ where E_r_ is the reduced modulus, E_s_ is Young’s modulus, v_s_ is Poisson’s ratio for a sample; E_i_=1140 GPa, and v_i_ is 0.07 for the diamond probe tip^43^.

### Atomic force microscopy (AFM) imaging

The biofilms (static medium) were grown on #1.5 round glass coverslips, 1 cm diameter (Ted Pella, Inc) in a 24-well plate. The biofilms were dried in air at room temperature (24-26°C) for 1 day before AFM imaging. Silicon AFM cantilevers (AC160TS-R3, k ≈ 30 N/m, Olympus) with a nominal tip radius of 7 nm were used with a scan angle of 90° and scan rates of 0.7 – 1.5 Hz. AFM in tapping mode was performed on Cypher (Oxford Instruments) at room temperature (23°C) in ambient air. The root-mean-square (r.m.s.) surface roughness of the biofilm over 2 *μ*m × 2 *μ*m areas was analyzed using built-in Cypher software (Oxford Instruments); these measurements were from the AFM height images of at least three biological replicates.

### AFM quantitative mapping of elasticity

The amyloid fibrils were induced by 250 ng/mL aTc at a diluted seeding density of 7.53 × 10^5^ cells/mL in the supplemented M63 medium at 30°C for 44 hours on polystyrene substrates with a Young’s modulus of 2.7 GPa (Bruker). The fibrils on the substrates were then washed twice with deionized water and dried at room temperature (24-26°C) for 1 day prior to the AFM characterization. Because of the nanoscale width of the CsgA amyloid fibrils, we used amplitude modulation-frequency modulation (AM-FM) AFM in tapping mode, which allows quantitative mapping of elasticity^44,45^. Amplitude modulation-frequency modulation (AM-FM) AFM in tapping mode was performed on Cypher (Oxford Instruments) at room temperature (23°C) in ambient air. The AM-FM AFM technology uses simultaneous monitoring of two distinct resonance modes of vibration of the cantilever^44,45^. Silicon AFM cantilevers (AC160TS-R3, k ≈ 30 N/m, Olympus) with a nominal tip radius of 7 nm were used with a scan angle of 90° and scan rates of 0.7 – 1.5 Hz. Prior to tip-sample interaction, the first cantilever drive frequency was adjusted to its resonant frequency (f_1_ ≈ 300 kHz) with a large free air amplitude, A_1_ = 2V, and the second frequency was tuned to its resonant frequency (f_2_ ≈ 1.6 MHz) with a much smaller free amplitude, A_2_ = 25 mV. For AFM tip calibration, a polystyrene film (Bruker) with the known modulus of 2.7 GPa was used as a substrate. The cantilever tip was calibrated to match the polystyrene modulus by adjusting a tip radius in the built-in Cypher software (Oxford Instruments). The average Young’s modulus of the amyloid fibrils was obtained assuming Poisson’s ratio of 0.3 by cross-section analysis of more than 70 fibrils from more than three AFM samples.

### AFM force spectroscopy

The bacterial cells were prepared on #1.5 round glass coverslips, 10 mm diameter (Ted Pella, Inc), by placing a 70 *μ*L seed density of 2.26 × 10^9^ cells/mL in deionized water. The cells were then dried at room temperature (24-26°C) for 2 days to evaporate solvent. AFM force spectroscopy in contact mode was conducted on MFP-3D (Oxford Instruments) at room temperature (23°C) in ambient air. Silicon nitride AFM cantilevers (TR800PSA, k ≈ 0.15 N/m, Olympus) with 200 *μ*m in length and a tip radius of 15 nm were used. To obtain Young’s modulus, the acquired retract force-distance curves were fitted with the Johnson-Kendall-Roberts (JKR) model^46^ assuming a Poisson’s ratio of 0.3, by using a software Scanning Probe Image Processor (SPIP, Image Metrology A/S). The average Young’s modulus was obtained from 16 measurements per sample for at least three AFM samples.

### SWNT dispersion, purification, and functionalization

SWNTs, 1 nm in diameter and 500 nm in length (HiPco Raw, Nanointegris), were dispersed in 2% w/v sodium cholate for 1 h at 15°C by an ultrasonic processor (Cole-Parmer). Purified SWNTs were then collected from supernatants after centrifugation at 32,000 r.p.m for 6 hours at 4°C. The SWNTs were quantified by measuring absorbance. The sodium cholate was removed and functionalized with SpyCatcher at the same time via dialysis based on a protein-SWNT complexation method^47^. SpyCatcher was added to SWNTs in 2% w/v sodium cholate to make a 1,000:1 ratio of SpyCatcher to SWNT. Then, SpyCatcher and SWNTs in sodium cholate were dialyzed via a 12-14 kDa cutoff membrane (Spectrum Laboratories) against three solutions for 3 days in total. The ionic strength of dialysis solutions was set to 10 mM NaCl for the first 2 days. For the first day, the pH was decreased to 5.4 by adding 0.1N HCl to make SpyCatcher (isoelectric point of 4.85) weakly acidic, in order to minimize the initial electrostatic repulsion between SpyCatcher and the SWNTs, thus facilitating the functionalization. After 1 day, the pH was adjusted to 11.0 by using 10N NaOH to increase the colloidal stability of the SWNTs functionalized with SpyCatcher. On the last day, phosphate buffered saline (PBS, pH7.4) was used as a final solution. The dialysis solutions were frequently replaced to increase purification efficiency. To remove unbound SpyCatcher, an Amicon filter (30kDa cutoff, EMD Millipore) was used with centrifugation at 7,000 r.p.m for 5 min.

### SWNT quantification

Based on the average diameter and length of SWNTs used in this study (1 nm and 500 nm, respectively), the number of carbon atoms per SWNT was calculated as 3.76 × 10^19^ C/m^2^ × 2 π × 0.5 nm × 500 nm = 5.903 × 10^4^ C, assuming graphene has an atomic density of 3.76 × 10^19^ C/m^2^. Thus, the molecular weight of the SWNT was calculated as 5.903 × 10^4^ × 12 g/mol = 7.0838 × 10^5^ g/mol. Inversion of the molecular weight yields 8.494 × 10^11^ /*μ*g. The SWNTs were dispersed in 2% m/v sodium cholate. The concentration of SWNTs can be calculated from absorbance (Abs) at 1cm-path length, based on a previous study^47^. For HipCo SWNTs, Abs_632nm_ × 27.8 gives *μ*g/mL. If Abs_632nm_ is 0.948, the SWNT concentration can be estimated as 0.948 × 27.8 = 26.35 *μ*g/mL. This concentration corresponds to 2.24 × 10^11^ /mL by 26.35 *μ*g/mL × 8.494 × 10^11^ /*μ*g.

### SpyCatcher expression and purification

The SpyCatcher proteins were heterologously expressed in an *E. coli* strain and purified (Supplementary Tables 1, 2). A DNA sequence encoding two cysteine and six histidine residues was placed at the N-terminus of SpyCatcher. The His residues were inserted for both nickel-affinity protein purification and SWNT functionalization. The *E. coli* strain was grown overnight in LB-Miller medium with appropriate antibiotics. Then, 10 mL stationary phase cells were added to 1 L LB-Miller to re-grow to OD600 of 0.5-0.7 at 37°C with shaking for 3-4 hours. Finally, IPTG (0.4 mM) was added and the culture was grown with shaking for 4 hours at 30°C. The cells were collected by centrifugation and lysed. Proteins were purified with Ni-NTA Resin (Qiagen) using standard protocols. 2 mM β-mercaptoethanol was used to break disulfide bonds. Following buffer exchange with Amicon columns, the proteins were resuspended in PBS. For further purification, SpyCatcher eluted in buffer (NPI-500, QIAGEN) was loaded into 0.5 mL Amicon filters (MWCO 3 kDa), PBS was added to the filters to make the total volume of 500 µL, and the samples were centrifuged at 11,000 rpm for 10 min. The washing with 400 *μ*L PBS and the centrifugation steps were repeated three times to remove imidazole from the protein solutions. The proteins were characterized by electrophoresis using a 12% SDS polyacrylamide gel and imaged using a ChemiDoc Touch Imaging System (Bio-Rad).

### SpyCatcher quantification

The concentration of SpyCatcher was calculated from Abs_280 nm_ at 1cm-path length. The molar extinction coefficient was 15,930 M^−1^cm^−1^ at Abs_280nm_ for SpyCatcher. Abs_280nm_ was divided by the molar extinction coefficient to yield the molar concentration.

### SWNT assembly on biofilms

To selectively bind the SWNTs functionalized with SpyCatcher on the SpyTag amyloid fibrils, SpyCatcher-SWNT was resuspended in PBS containing 0.05% v/v Tween-20 and 0.1% w/v bovine serum albumin (BSA). Solutions having a 1000:1 SpyCatcher-to-SWNT ratio at given concentrations were incubated on the SpyTag biofilms for 2 hours at room temperature (24-26°C) with slow rocking. The biofilms were then washed once with the PBS-Tween-BSA buffer and three times with deionized water. For instrumented indentation, the SWNT-integrated biofilms were dried in air at room temperature (24-26°C) for 1 day. For transmission electron microscope (TEM) analysis, the SWNT-assembled biofilms were scraped by pipetting and placed on a lacey carbon TEM grid with copper (Ted Pella, Inc.) for 20 min. The amyloid fibrils were stained with 2% uranyl acetate for 10 seconds, washed twice with deionized water, and dried in air for an hour. The images were recorded using a Tecnai G2 Spirit Twin TEM (FEI).

### Metal-sensing biofilms

The biofilms (static medium) were grown in a 96-well plate. The biofilms were then incubated at room temperature (24-26°C) in a M63 medium supplemented with 0.2% w/v glucose and 1 mM MgSO_4_ containing given Cd^2+^ concentrations. The optical density at 600 nm (OD_600_) and GFP fluorescence signals from the biofilms were recorded with excitation at 485 nm and emission at 515 nm by a multi-mode microplate reader (Synergy H1, BioTek Instruments, Inc.) using a bottom detector and a sensitivity of 100. In the kinetic response of the biofilms, to offset fluorescence signals from planktonic cells in the M63 medium, the relative fluorescence units (RFU) were normalized to OD_600_ of the biofilms in the M63 medium and reported as RFU/OD_600_. In the end-point measurements, the biofilms were incubated at room temperature for 9 hours in the M63 medium. To detect fluorescence signals emitted only from the biofilms, planktonic cells were removed by replacing the existing M63 medium with fresh M63 medium. For colony-forming unit (CFU) counts, the biofilms were disrupted by pipetting thoroughly and homogenized with a stainless bead beater 5 mm in diameter (Qiagen) for 10 min at a frequency of 30 beats per second by TissueLyser II (Qiagen). The recorded fluorescence intensities were normalized to CFU/cm^2^ and reported in RFU/CFU.

### Statistics

Statistical analysis was carried out using GraphPad Prism (GraphPad Software). Data are presented as mean ± s.d. (standard deviation). The s.d. values were obtained from at least three independent samples. Box-plots are presented as center lines (median), box limits (upper and lower quartiles), whiskers (1.5× interquartile range), and points (outliers) for n > 10 from at least three independent samples. Statistical significance between groups was determined using an unpaired, two-tailed Student’s t-test assuming unequal s.d.

## Data availability

All data supporting the findings of this study are available within the article and its supplementary information or from the corresponding author upon request.

## Acknowledgements

H.P. thanks Dr. Karen Pepper for editing the manuscript and Dr. Jiyoung Ahn in the Strano group at MIT for helping with the SWNT photoluminescence measurement. T.K.L. gratefully acknowledges funding for this study received from the Army Research Office (Awards no. W911NF-11-1-0281 and W911NF-17-2-0077), the Institute for Soldier Nanotechnologies/Army (Award no. W9111NF-13-D-0001), the NIH (Award no. 18047174 and, via the CISB - MIT Center for Integrative Synthetic Biology, Award no. 1-P50GM098792-01A1), the NSF (Award no. CCF-1521925), the Defense Threat Reduction Agency (Awards no. HDTRA1-14-1-0007 and HDTRA1-15-1-0050), DARPA (Award no. HR0011-15-C-0084 and, via Harvard Medical School, Award no. 17126066), and The Leona M. and Harry B. Helmsley Charitable Trust (Award no. 3239). T.-C.T thanks the J-WAFS Graduate Student Fellowship for its financial support.

## Author contributions

H.P. and T.K.L. conceived the research. T.K.L. supervised the research. H.P. designed and performed the experiments. A.S. helped H.P. with discussions and assisted H.P. in measuring mechanical properties by instrumented indentation and AFM. T.-C.T. helped H.P. with the metal-sensing experiment. L.W. helped H.P. with the microfluidic chamber design and fabrication. H.P. and T.K.L. analyzed the data and wrote the manuscript with help from all authors.

## Competing interests

T.K.L. is a co-founder of Senti Biosciences, Synlogic, Engine Biosciences, Tango Therapeutics, Corvium, BiomX, Eligo Biosciences, Bota.Bio, Avendesora, and NE47Bio. T.K.L. also holds financial interests in nest.bio, Armata, IndieBio, MedicusTek, Quark Biosciences, Personal Genomics, Thryve, Lexent Bio, MitoLab, Vulcan, Serotiny, Avendesora, Pulmobiotics, Provectus Algae, Invaio, and NSG Biolabs.

Other authors declare no competing interests.

## Additional information

**Supplementary information** is available for this paper.

Correspondence and requests for materials should be addressed to T.K.L.

## References

1. Wegst, U. G. K. & Ashby, M. F. The mechanical efficiency of natural materials. Philos. Mag. (2004). doi:10.1080/14786430410001680935

2. Fratzl, P. & Weinkamer, R. Nature’s hierarchical materials. Progress in Materials Science (2007). doi:10.1016/j.pmatsci.2007.06.001

3. Wegst, U. G. K., Bai, H., Saiz, E., Tomsia, A. P. & Ritchie, R. O. Bioinspired structural materials. Nat. Mater. 14, 23–36 (2015).

4. Flemming, H. C. et al. Biofilms: An emergent form of bacterial life. Nature Reviews Microbiology (2016). doi:10.1038/nrmicro.2016.94

5. Vidakovic, L., Singh, P. K., Hartmann, R., Nadell, C. D. & Drescher, K. Dynamic biofilm architecture confers individual and collective mechanisms of viral protection. Nat. Microbiol. (2017). doi:10.1038/s41564-017-0050-1

6. Flemming, H. C. & Wuertz, S. Bacteria and archaea on Earth and their abundance in biofilms. Nat. Rev. Microbiol. (2019). doi:10.1038/s41579-019-0158-9

7. Persat, A. et al. The mechanical world of bacteria. Cell (2015). doi:10.1016/j.cell.2015.05.005

8. Hobley, L., Harkins, C., MacPhee, C. E. & Stanley-Wall, N. R. Giving structure to the biofilm matrix: An overview of individual strategies and emerging common themes. FEMS Microbiology Reviews (2015). doi:10.1093/femsre/fuv015

9. Hall-Stoodley, L. & Stoodley, P. Evolving concepts in biofilm infections. Cellular Microbiology (2009). doi:10.1111/j.1462-5822.2009.01323.x

10. Battin, T. J., Kaplan, L. A., Newbold, J. D. & Hansen, C. M. E. Contributions of microbial biofilms to ecosystem processes in stream mesocosms. Nature (2003). doi:10.1038/nature02152

11. Johnsen, A. R. & Karlson, U. Evaluation of bacterial strategies to promote the bioavailability of polycyclic aromatic hydrocarbons. Appl. Microbiol. Biotechnol. (2004). doi:10.1007/s00253-003-1265-z

12. Fuchs, S., Haritopoulou, T., Schäfer, M. & Wilhelmi, M. Heavy metals in freshwater ecosystems introduced by urban rainwater runoff - Monitoring of suspended solids, river sediments and biofilms. in Water Science and Technology (1997). doi:10.1016/S0273-1223(97)00586-6

13. Halan, B., Buehler, K. & Schmid, A. Biofilms as living catalysts in continuous chemical syntheses. Trends in Biotechnology (2012). doi:10.1016/j.tibtech.2012.05.003

14. Stoodley, P., Lewandowski, Z., Boyle, J. D. & Lappin-Scott, H. M. Structural deformation of bacterial biofilms caused by short-term fluctuations in fluid shear: An in situ investigation of biofilm rheology. Biotechnol. Bioeng. (1999). doi:10.1002/(SICI)1097-0290(19991005)65:1<83::AID-BIT10>3.0.CO;2-B

15. Koza, A., Hallett, P. D., Moon, C. D. & Spiers, A. J. Characterization of a novel air-liquid interface biofilm of Pseudomonas fluorescens SBW25. Microbiology (2009). doi:10.1099/mic.0.025064-0

16. Jones, W. L., Sutton, M. P., Mckittrick, L. & Stewart, P. S. Chemical and antimicrobial treatments change the viscoelastic properties of bacterial biofilms. Biofouling (2011). doi:10.1080/08927014.2011.554977

17. Wu, C., Lim, J. Y., Fuller, G. G. & Cegelski, L. Quantitative analysis of amyloid-integrated biofilms formed by uropathogenic escherichia coli at the air-liquid interface. Biophys. J. (2012). doi:10.1016/j.bpj.2012.06.049

18. Chen, A. Y. et al. Synthesis and patterning of tunable multiscale materials with engineered cells. Nat. Mater. 13, 515–23 (2014).

19. Nguyen, P. Q., Botyanszki, Z., Tay, P. K. R. & Joshi, N. S. Programmable biofilm-based materials from engineered curli nanofibres. Nat. Commun. (2014). doi:10.1038/ncomms5945

20. Huang, J. et al. Programmable and printable Bacillus subtilis biofilms as engineered living materials. Nat. Chem. Biol. (2019). doi:10.1038/s41589-018-0169-2

21. Vidal, O. et al. Isolation of an Escherichia coli K-12 mutant strain able to form biofilms on inert surfaces: Involvement of a new ompR allele that increases curli expression. J. Bacteriol. (1998).

22. Prigent-Combaret, C. et al. Developmental pathway for biofilm formation in curli-producing Escherichia coli strains: Role of flagella, curli and colanic acid. Environ. Microbiol. (2000). doi:10.1046/j.1462-2920.2000.00128.x

23. Oliver, W. C. & Pharr, G. M. An improved technique for determining hardness and elastic modulus (Young’s modulus). J. Mater. Res. (1992).

24. Körstgens, V., Flemming, H. C., Wingender, J. & Borchard, W. Uniaxial compression measurement device for investigation of the mechanical stability of biofilms. J. Microbiol. Methods (2001). doi:10.1016/S0167-7012(01)00248-2

25. Baniasadi, M. et al. Nanoindentation of Pseudomonas aeruginosa bacterial biofilm using atomic force microscopy. Mater. Res. Express (2015). doi:10.1088/2053-1591/1/4/045411

26. Kundukad, B. et al. Mechanical properties of the superficial biofilm layer determine the architecture of biofilms. Soft Matter (2016). doi:10.1039/c6sm00687f

27. Kesel, S. et al. Direct comparison of physical properties of Bacillus subtilis NCIB 3610 and B-1 biofilms. Appl. Environ. Microbiol. (2016). doi:10.1128/AEM.03957-15

28. Asally, M. et al. Localized cell death focuses mechanical forces during 3D patterning in a biofilm. Proc. Natl. Acad. Sci. U. S. A. (2012). doi:10.1073/pnas.1212429109

29. Heydorn, A. et al. Quantification of biofilm structures by the novel computer program COMSTAT. Microbiology (2000). doi:10.1099/00221287-146-10-2395

30. Knowles, T. P. J. & Buehler, M. J. Nanomechanics of functional and pathological amyloid materials. Nature Nanotechnology (2011). doi:10.1038/nnano.2011.102

31. Knowles, T. P. et al. Role of intermolecular forces in defining material properties of protein nanofibrils. Science (80-.). (2007). doi:10.1126/science.1150057

32. Chapman, M. R. et al. Role of Escherichia coli curli operons in directing amyloid fiber formation. Science (80-.). (2002). doi:10.1126/science.1067484

33. Treacy, M. M. J., Ebbesen, T. W. & Gibson, J. M. Exceptionally high Young’s modulus observed for individual carbon nanotubes. Nature (1996). doi:10.1038/381678a0

34. Zakeri, B. et al. Peptide tag forming a rapid covalent bond to a protein, through engineering a bacterial adhesin. Proc. Natl. Acad. Sci. U. S. A. (2012). doi:10.1073/pnas.1115485109

35. Wang, S. et al. Peptides with selective affinity for carbon nanotubes. Nat. Mater. (2003). doi:10.1038/nmat833

36. O’Connell, M. J. et al. Band gap fluorescence from individual single-walled carbon nanotubes. Science (80-.). (2002). doi:10.1126/science.1072631

37. Tang, T. C. et al. Hydrogel-based biocontainment of bacteria for continuous sensing and computation. Nat. Chem. Biol. (2021). doi:10.1038/s41589-021-00779-6

38. Colvin, K. M. et al. The Pel and Psl polysaccharides provide Pseudomonas aeruginosa structural redundancy within the biofilm matrix. Environ. Microbiol. (2012). doi:10.1111/j.1462-2920.2011.02657.x

39. Hobley, L. et al. BslA is a self-assembling bacterial hydrophobin that coats the Bacillus subtilis biofilm. Proc. Natl. Acad. Sci. 110, 13600–13605 (2013).

40. Gilbert, C. et al. Living materials with programmable functionalities grown from engineered microbial co-cultures. Nat. Mater. (2021). doi:10.1038/s41563-020-00857-5

41. Moser, F., Tham, E., González, L. M., Lu, T. K. & Voigt, C. A. Light-Controlled, High-Resolution Patterning of Living Engineered Bacteria Onto Textiles, Ceramics, and Plastic. Adv. Funct. Mater. (2019). doi:10.1002/adfm.201901788

42. Wang, L. et al. Construction of oxygen and chemical concentration gradients in a single microfluidic device for studying tumor cell-drug interactions in a dynamic hypoxia microenvironment. Lab Chip (2013). doi:10.1039/c2lc40661f

43. Johnson, K. L. Contact Mechanics. (1989). doi:10.1201/b17110-2

44. Kocun, M., Labuda, A., Meinhold, W., Revenko, I. & Proksch, R. Fast, High Resolution, and Wide Modulus Range Nanomechanical Mapping with Bimodal Tapping Mode. ACS Nano (2017). doi:10.1021/acsnano.7b04530

45. Benaglia, S., Gisbert, V. G., Perrino, A. P., Amo, C. A. & Garcia, R. Fast and high-resolution mapping of elastic properties of biomolecules and polymers with bimodal AFM. Nat. Protoc. (2018). doi:10.1038/s41596-018-0070-1

46. Efremov, Y. M., Bagrov, D. V., Kirpichnikov, M. P. & Shaitan, K. V. Application of the Johnson-Kendall-Roberts model in AFM-based mechanical measurements on cells and gel. Colloids Surfaces B Biointerfaces (2015). doi:10.1016/j.colsurfb.2015.06.044

47. Dang, X. et al. Virus-templated self-assembled single-walled carbon nanotubes for highly efficient electron collection in photovoltaic devices. Nat. Nanotechnol. (2011). doi:10.1038/nnano.2011.50

