## Supplementary Figures 1-9 and Tables 1-2 for "Ultra-lightweight living structural material for enhanced stiffness and environmental sensing"

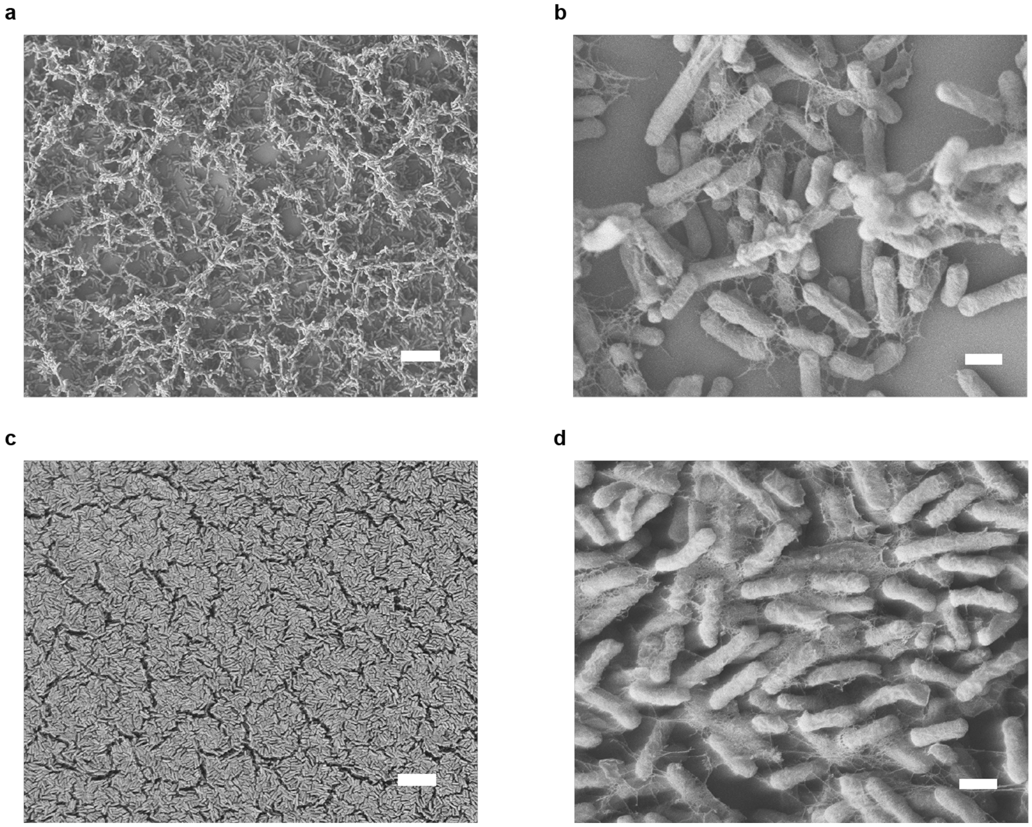


**Supplementary Fig. 1. Scanning electron microscopy (SEM) images of biofilms. a**, **b**, A porous structure was revealed when a diluted seeding density of $5.65 \times 10^7$ cells/mL was used to grow biofilms. **c**, **d**, A closely packed structure of the biofilm surface formed when seeding density was increased to $2.26 \times 10^8$ cells/mL for biofilm growth. **a**, **c**, Scale bar, 10 µm. **b**, **d**, Scale bar, 1 µm.

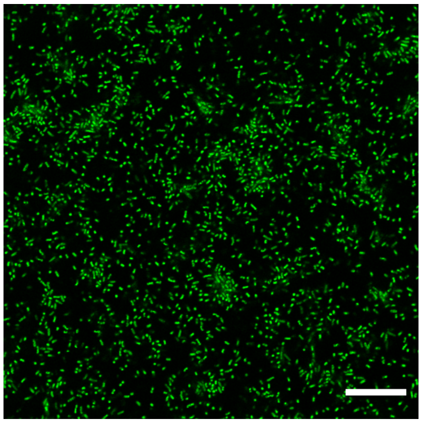

**Supplementary Fig. 2. Confocal microscopy image of the amyloid fibrils in the densely packed biofilm.** The production of amyloid fibrils by *E. coli* ompR234 cells was confirmed by staining with thioflavin T. Scale bar, 20 µm.

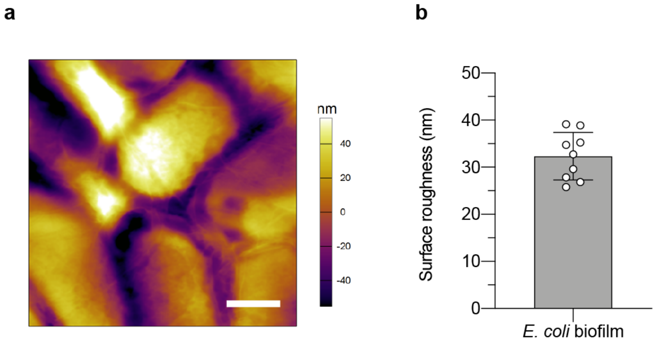


**Supplementary Fig. 3. a,** Atomic force microscopy (AFM) height image over the static medium biofilm surface. Scale bar, 400 nm. **b**, The root-mean-square surface roughness of the biofilm over 2 *μ*m × 2 *μ*m areas was determined by analysis of AFM height images. Data are mean ± s.d. for n = 9 from three independent samples.

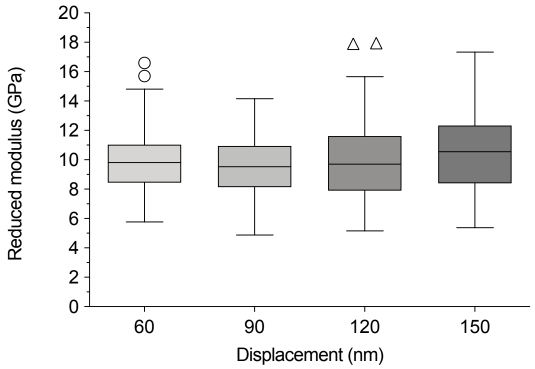


**Supplementary Fig. 4. Results of instrumented indentation on the biofilms.** Reduced elastic moduli of 60, 90, 120, and 150 nm indenting depths analyzed from the load-displacement curves for the static medium biofilms. Box-plots show center lines (median), box limits (upper and lower quartiles), whiskers (1.5× interquartile range), and points (outliers) for n ≥ 140 from three independent samples.

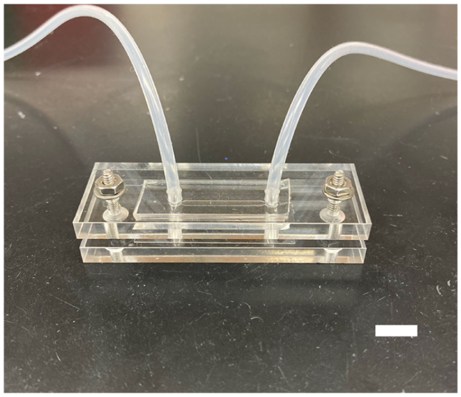

**Supplementary Fig. 5. Fully assembled microfluidic chamber device.** The dimensions of a single-channel PDMS flow chamber were 250 $\mu$m high, 2 mm wide, and 20 mm long. Scale bar, 1 cm.

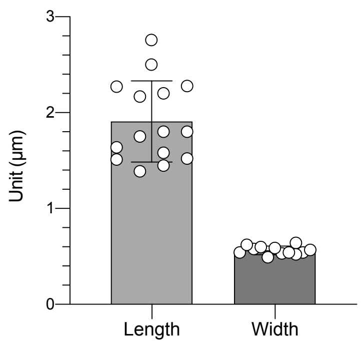


**Supplementary Fig. 6. Dimensions of the *E. coli* ompR234 cells.** The length and width of the cells were estimated from SEM images. Data are mean ± s.d. for n = 16 for length and n = 13 for width from three independent samples.

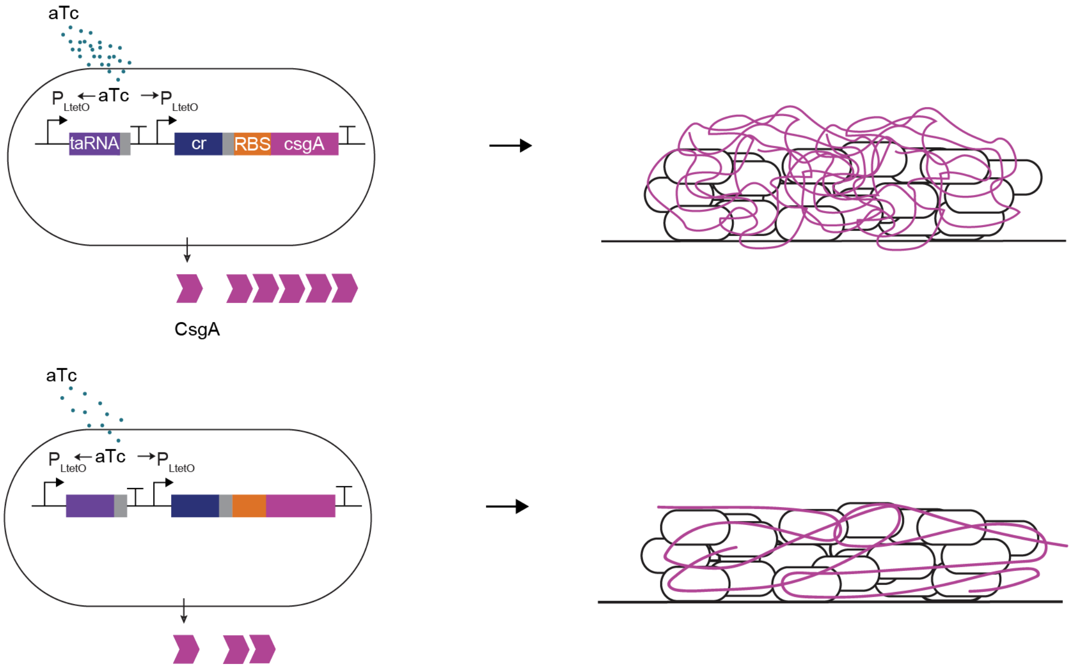


**Supplementary Fig. 7. Schematic of the dose effect of the lower concentrations of the inducer, aTc.** A lower concentration of aTc produces a smaller quantity of fibrils (bottom panel), whereas a higher concentration of aTc induces a larger quantity of fibrils (upper panel).

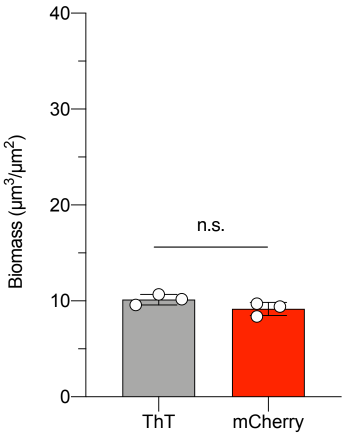


**Supplementary Fig. 8. Comparison between biomasses of amyloid fibrils and *E. coli* cells.** In the same biofilms, the biomass of the amyloid fibrils stained with thioflavin T (ThT) was not significantly different from the biomass of the cells constitutively expressing mCherry. Thus, the biomass analyzed from the mCherry-expressing cells also indicates the approximate biomass of the amyloid fibrils. Not significant (n.s.) $P > 0.05$, Student t test. Data are mean ± s.d. for $n = 3$ independent samples.

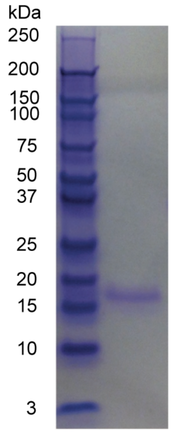


**Supplementary Fig. 9. Gel image of SpyCatcher.** A polyacrylamide gel stained with Coomassie confirms the presence of SpyCatcher, a 16-kDa protein.

**Supplementary Table 1.** Bacterial strains used in this study.

| Strain name | Strain ID | Description | Antibiotic Resistance | Source |
|---|---|---|---|---|
| MG1655 *PRO* Δ*csgA* *ompR234* | fAYC002 | *E. coli* host strain with *PRO* cassette ($P_{laci}^{q}$/*lacI*, $P_{N25}$/*tetR*, $Spec^{R}$) that constitutively expresses TetR and LacI repressors, with knock-out of endogenous *csgA*, and with an *ompR234* allele that confers the ability to produce fibrils in liquid M63 minimal media. | Spec, Kan | [1] |
| aTc/CsgA | fAYC008 | *E. coli* strain that expresses CsgA under tight regulation by an anhydrotetracycline (aTc) inducer-responsive riboregulator. Made by transforming pZA-CmR-rr12-pL(tetO)-*csgA* plasmid into MG1655 *PRO* Δ*csgA ompR234*. | Spec, Kan, Cm | [2] |
| aTc/$CsgA_{SpyTag}$ | fAYC024 | *E. coli* strain that expresses $CsgA_{SpyTag}$ under tight regulation by an aTc inducer-responsive riboregulator. Made by transforming pZA-CmR-rr12-pL(tetO)-$csgA_{SpyTag}$ into MG1655 *PRO ΔcsgA ompR234.* | Spec, Kan, Cm | This study |
| Cd,Pb,Zn/GFP + aTc/CsgA | FHE002 | *E. coli* strain that expresses CsgA under tight regulation by an aTc inducer-responsive riboregulatory, and expresses GFP upon detection of Cd, Pb, Zn ions. Made by co-transforming $P_{zntA}$-*gfp* and pZA-CmR-rr12-pL(tetO)-*csgA* into MG1655 PRO *ΔcsgA ompR234*. | Spec, Kan, Cm, Amp | This study |
| mCherry + aTc/$CsgA_{His}$ | fAYC011 | *E. coli* strain that expresses CsgA under tight regulation by an aTc inducer-responsive riboregulatory, and constitutively expresses mCherry. Made by co-transforming pZE-AmpR_proB-mCherry and pZA-CmR-rr12-pL(tetO)-$csgA_{His}$ into MG1655 *PRO ΔcsgA ompR234.* | Spec, Kan, Cm, Amp | [2] |
| BL21(DE3) pLysS / pDEST14-T7-*Cys₂-His₆-SpyCatcher* | fAYC016 | *E. coli* strain that expresses $His_2$-$Cys_2$-SpyCatcher when induced by IPTG. Made by transforming pDEST14-T7-$Cys_2$-$His_6$-*SpyCatcher* into BL21(DE3) pLysS. | Cm, Amp | [2] |

**Supplementary Table 2.** Plasmids used in this study.

| Plasmid name | Plasmid ID | Description | Source |
|---|---|---|---|
| pZA-CmR-rr12-pL(tetO)-*csgA* | pAYC002 | p15A origin, Cm resistance, rr12 riboregulator, pL(tetO) promoter, *csgA* output gene | [2] |
| pZA-CmR-rr12-pL(tetO)-$csgA_{SpyTag}$ | pAYC021 | p15A origin, Cm resistance, rr12 riboregulator, pL(tetO) promoter, $csgA_{SpyTag}$ output gene | This study |
| $P_{zntA}$-*gfp* | pEZ074 | $P_{zntA}$ promoter, *gfp* output gene | [3] |
| pZE-AmpR-proB-*mCherry* | pAYC010 | ColE1 origin, Amp resistance, proB promoter, *mCherry* output gene | [2] |
| pZA-CmR-rr12y-pLuxR-$csgA_{His}$ | pAYC007 | p15A origin, Cm resistance, rr12 riboregulator, pL(tetO) promoter, $csgA_{His}$ output gene | [2] |
| pDEST14-T7-$Cys_2$-$His_6$-*SpyCatcher* | pAYC016 | pBR322 origin, Amp resistance, T7 promoter, $Cys_2$-$His_6$-*SpyCatcher* output gene | [2] |

**References**


1. Prigent-Combaret, C. *et al.* Complex regulatory network controls initial adhesion and biofilm formation in Escherichia coli via regulation of the csgD gene. *J. Bacteriol.* (2001). doi:10.1128/JB.183.24.7213-7223.2001
2. Chen, A. Y. *et al.* Synthesis and patterning of tunable multiscale materials with engineered cells. *Nat. Mater.* **13**, 515–23 (2014).
3. Tang, T. C. *et al.* Hydrogel-based biocontainment of bacteria for continuous sensing and computation. *Nat. Chem. Biol.* (2021). doi:10.1038/s41589-021-00779-6
